# Co-localization of antibiotic resistance genes is widespread in the infant gut microbiome and associates with an immature gut microbial composition

**DOI:** 10.1101/2023.02.19.525876

**Authors:** Xuanji Li, Asker Brejnrod, Urvish Trivedi, Jakob Russel, Jakob Stokholm, Jonathan Thorsen, Shiraz A Shah, Gisle Alberg Vestergaard, Morten Arendt Rasmussen, Joseph Nesme, Hans Bisgaard, Søren Johannes Sørensen

## Abstract

Even in the absence of antibiotic exposure, the gut microbiome of infants has been shown to contain numerous antibiotic resistance genes (ARGs), but the mechanism for this remains unclear. In environmental bacteria, the selective advantage of ARGs can be increased through co-localization with genes such as other ARGs, biocide resistance genes, metal resistance genes, and virulence genes. However, this phenomenon is unknown from the human gut microbiome during early life despite frequent exposures to biocides and metals from an early age. Here, we conducted a comprehensive analysis of genetic co-localization of resistance genes in a cohort of 662 Danish children and examined the association between such co-localization and environmental factors as well as gut microbial maturation. Our study showed that co-localization of ARGs with other resistance and virulence genes is common in the early gut microbiome and is associated with gut bacteria that are indicative of low maturity, especially *E. coli*. The most common forms of co-localization involved tetracycline and fluoroquinolone resistance genes located near other ARGs, and, on plasmids, co-localization predominantly occurred in the form of class 1 integrons. Antibiotic use caused a short-term increase in mobile ARGs, while non-mobile ARGs showed no significant change. Finally, we found that a high abundance of virulence genes was associated with low gut microbial maturity and that virulence genes showed even higher potential for mobility than ARGs. Our study highlights important constraints that need to be considered when developing strategies to combat ARG dissemination.

## Main

The first years of life are pivotal for maturation of the gut microbiome ^1,2^ and healthy development of the host immune system ^3^. Prolonged perturbation of the developing gut microbiome is linked to an increased risk for subsequent diseases, such as asthma ^4^ and allergies ^5^. Together with the maturing gut microbiome, the antibiotic resistome—the pool of genes that contribute to antimicrobial resistance—also develops in the infant gut in the first few years of life. We recently showed that enrichment in antibiotic resistance genes (ARGs) in early life was associated with gut microbial immaturity, similar to an asthma-associated microbiome composition ^6^. The enrichment of ARGs was mainly driven by the presence of *E. coli* in the infant gut ^6^. However, the most prevalent ARGs in the infant gut encoded resistance against antibiotics that neither the children nor their mothers had taken ^6^. We thus speculate that the maintenance of ARGs in gut bacteria occurs partly through a mechanism of “co-selection” between ARGs and other resistance genes. The physical linkage of multiple genes encoding different resistance phenotypes on the same genetic element (co-localization) is a widespread phenomenon ^7^ and because of this, selection for resistance against one antibacterial agent can result in the maintenance of resistance against other agents, termed “co-selection”^8^ (Fig. 1). Co-selection can favor a variety of ARGs in bacterial hosts that are more likely to be exposed to diverse co-selection agents. Additionally, this kind of multi-agent resistance can further increase the persistence of certain bacteria and plasmids in the gut. The prolonged persistence of early colonizers such as *E. coli* can perturb subsequent colonization by beneficial commensals, thereby resulting in a delay in the microbial maturation of the infant gut.

**Fig 1.**
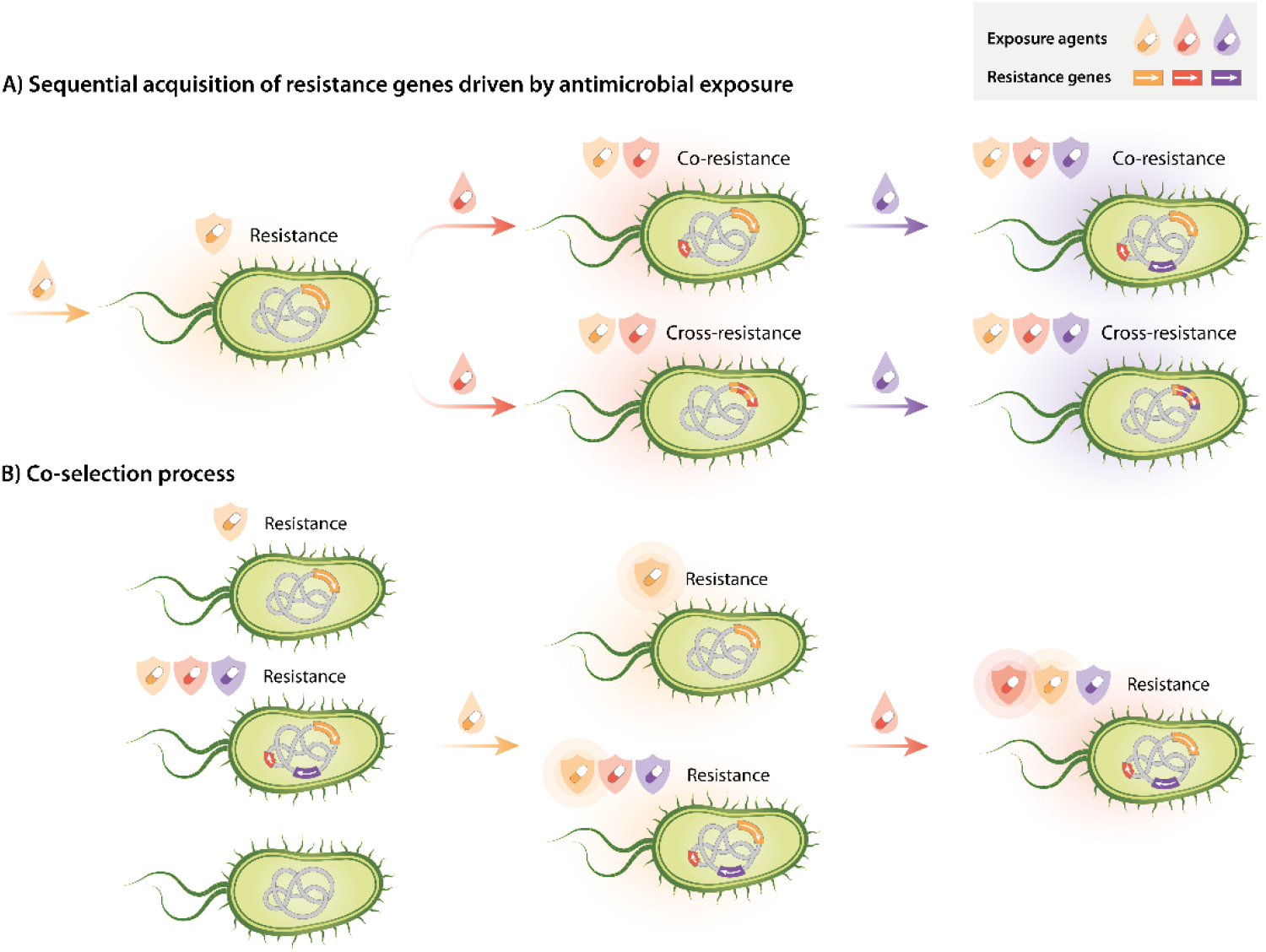
Emergence and co-selection of multidrug resistance in chromosomes or plasmids. **A)** Progressive acquisition of multiple resistances from mutations or gene transfer as a result of exposure to antimicrobial agents. Co-resistance represents the co-localization of multiple resistance genes in the same gene fragment. Cross-resistance represents a gene that confers resistance to multiple antimicrobials. **B)** Co-selection between multiple resistance genes upon exposure to any of the corresponding antimicrobials. This figure is adapted from the work of Cantón and Ruiz-Garbajosa ^8^.

Co-selection between ARGs and genes conferring resistance against agents such as antibacterial biocides and metals is frequently detected in the environment ^9^. Furthermore, these resistance genes are often located next to mobile genetic elements (MGEs) or on plasmids which facilitates their transfer between chromosomes and plasmids or between cells. However, the prevalence of this phenomenon in the human gut is largely unknown— especially in the developing infant gut. This is a critical knowledge gap considering that humans are frequently exposed to co-selective agents such as biocides and metals in their daily lives. Biocides are usually utilized as antiseptics on the skin to prevent microbial infection ^10^. Certain essential metals are indispensable for humans while some heavy metals can inadvertently enter the food chain ^11^. In addition, metals have been used against microbial infections in human and veterinary medicine for decades ^12^.

In addition to resistance genes, bacteria have numerous other genetic tools to help them persist and thrive in their environments. In particular, virulence genes play a critical role in determining bacterial pathogenicity and pose an important threat to human health ^13^; such genes typically enhance pathogenicity via modifications to a bacterium’s toxicity and/or invasiveness ^14^. Virulence genes have evolved with their bacterial hosts for millions of years ^14^, and like ARGs, many have been transferred among bacteria by horizontal gene transfer (HGT)^15,16^, which is one of the main ways in which human opportunistic pathogens acquire virulence in the course of evolution ^17^. However, the early establishment of virulence genes in the gut microbiota remains unclear. It is possible that co-localization between virulence genes and ARGs may confer a selective advantage on virulence genes in bacteria that are likely to be exposed to antibiotics; the overall result would be to improve the persistence of such bacteria in the infant gut. To date, though, there have been no comprehensive investigations on the co-localization of virulence genes and ARGs in the infant gut microbiome. A better understanding of co-selection phenomena during early life could help elucidate the maintenance and proliferation of both ARGs and virulence genes, and explore the main drivers for the persistence of gut bacteria in early life, which is of key importance for efforts to alleviate the spread of ARGs and virulence genes.

In this paper, we present a systematic analysis of co-localization between ARGs, metal and biocide resistance genes, and virulence genes. We investigate the presence of co-localized genes in MGEs and on plasmids and characterize the common bacterial hosts in which this phenomenon occurs. To do this, we used data obtained from metagenomic sequencing of 662 fecal samples from healthy infants in the Copenhagen Prospective Studies on Asthma in Childhood 2010 (COPSAC2010) birth cohort. We evaluated the effects of antibiotic use on mobile ARGs and investigated the relationship between co-localization and microbial maturation of the infant gut, particularly with regard to bacterial community profiles that have been linked to asthma.

## Results

### Co-localization in the infant gut was most common between tetracycline or fluoroquinolone ARGs and other ARGs

We previously showed that genes conferring resistance to fluoroquinolone and tetracycline were the most prevalent ARGs in the infant gut despite the fact that none of the infants in the study population had ever taken either of these drugs ^6^. In the absence of antibiotic exposure, we hypothesized that gut bacteria could maintain the acquired antimicrobial resistance by co-selection. We thus conducted a comprehensive investigation into the co-selection of ARGs, with a focus on two phenomena: 1) co-resistance (multiple ARGs conferring resistance to different drugs that are all located in the same genetic element (contig)), and 2) cross-resistance (multidrug resistant ARGs, MDR ARGs). Of all the contigs that were found to contain ARGs, 21.2% carried multiple ARGs and more than half of these (12.5% of all contigs) contained ARGs known to confer resistance against different drugs (Fig. 2A). Using metagenomically assembled genomes (MAGs), we traced the contigs carrying multiple ARGs with different resistance profiles to 55 bacterial species of origin, representing 5 phyla. The overwhelming majority (89%) of the traceable contigs were from Proteobacteria, particularly *E. coli* (Fig. 2B).

**Fig 2.**
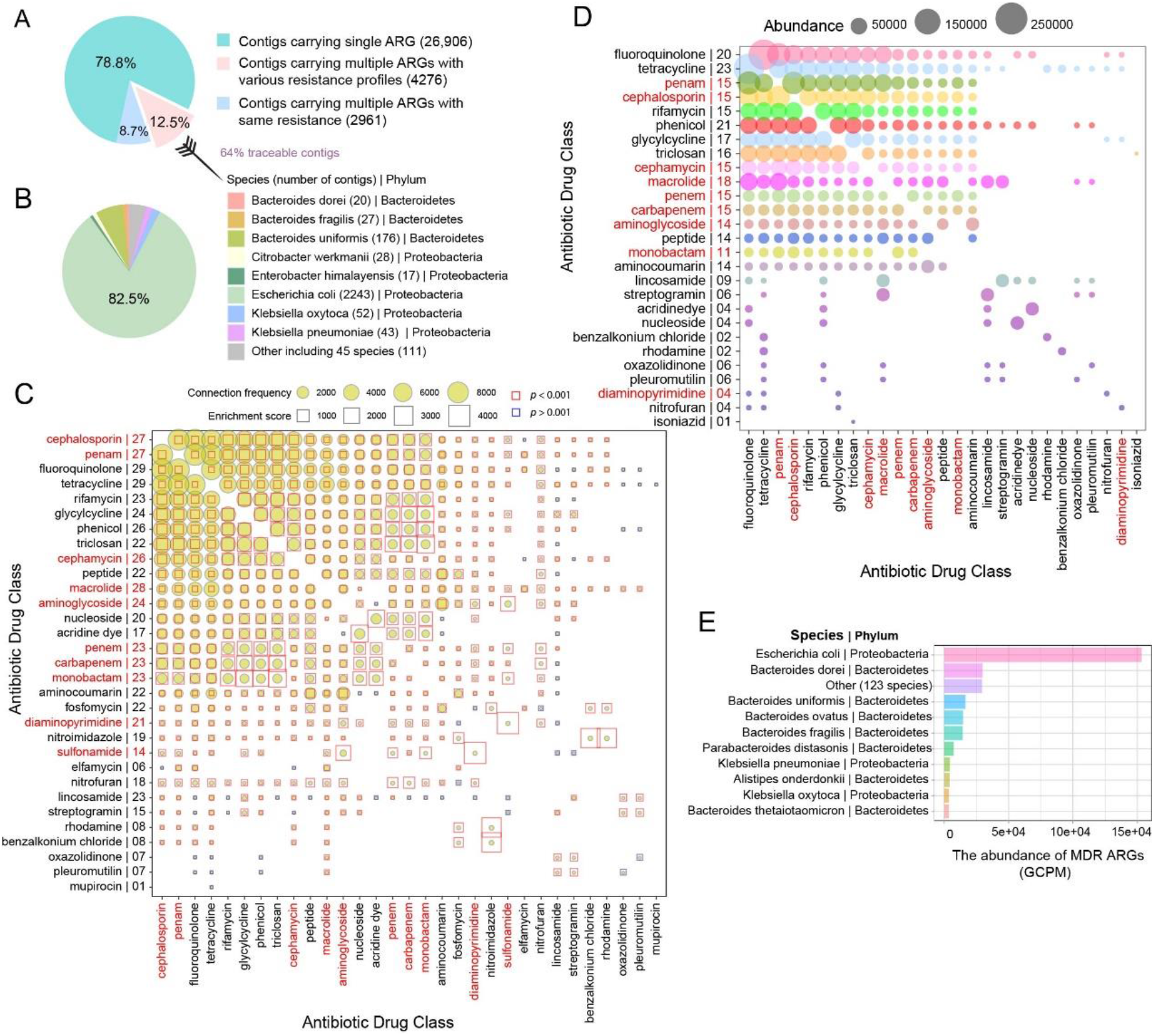
Overview of co-selection between ARGs in the infant gut. **A)** Proportion of contigs carrying different numbers of ARGs. **B)** Taxonomic origin (bacterial species and phylum) of contigs carrying multiple ARGs with different resistance profiles. **C)** Co-localization bubble chart representing the drug classes related to different ARGs. The frequency of connections between ARGs targeting different drug classes in the contigs, and the associated enrichment scores, are shown in the figure. On the y-axis, the number to the right of the name indicates the number of other drug classes represented in the co-localization arrangements; the drug classes in red were those taken by members of this cohort. The size of the bubble indicates the frequency of connections in the contigs. An enrichment score was calculated as the actual number of co-localization contigs/expected number of co-localization contigs with a given arrangement; scores higher than 1 are indicative of enrichment in that co-localization arrangement. A binomial test with FDR adjustment was used to test the statistical significance of enrichment patterns (*p* < 0.001 was set as significance cutoff; red square frame represents *p* < 0.001 and blue square frame represents *p* > 0.001). A significant *p*-value indicates that the occurrence of that specific pattern of gene co-localization would not be expected by chance. The size of the square frame represents the magnitude of the enrichment score. **D)** Co-localization bubble chart representing the drug classes targeted by 167 MDR ARGs. The size of the bubble is proportional to the abundance of MDR ARGs. On the y-axis, the number to the right of the name indicates the number of other drug classes represented in the co-localization arrangements. The drug classes in red were those taken by members of this cohort. **E)** The abundance of MDR ARGs in different bacteria at the phylum and species level.

To reflect gene co-resistance more intuitively, we transformed the co-resistance network of 4276 ARG-carrying contigs into a network that depicts resistance against 31 classes of drugs. We collapsed together all genes conferring resistance to a given drug class and examined how often a resistance gene for one class was found to be co-localized with a resistance gene for another class (Fig. 2C). Genes conferring resistance to cephalosporin, penam, fluoroquinolone, and tetracycline were the most likely to be co-localized with other drug resistance genes. Co-localization with fluoroquinolone and tetracycline resistance genes was most common, followed by macrolides, cephalosporin, and penam. Of these, cephalosporin, penam, and macrolides were commonly prescribed antibiotics for infants in this cohort. In addition, we assessed the stochasticity of co-localization using an enrichment score, which we defined as the fold difference between the actual and the expected numbers of co-localized contigs. An enrichment score higher than one indicated that a certain type of co-localization was enriched in the gut. Among different types of ARGs, enrichment scores for co-localization ranged from 1.65 to 4,126 (mean (SD): 365(554), Fig. 2C). We next used a binomial test to determine whether the number of contigs carrying multiple ARGs was significantly higher than the value expected by chance, i.e. whether the enrichment scores were statistically significant. With the exception of ARGs related to lincosamide, most of the co-localization between ARGs was significantly enriched compared to the expected values (binomial test; adjusted *p* < 0.001), suggesting that the co-localization was not the result of chance.

Among the 409 antibiotic resistance genes detected in the infant gut, 167 provided resistance against at least two antibiotics. These multidrug resistant (MDR) ARGs conferred resistance against 27 drug classes in total. Of these, cross-resistance to tetracycline, phenicol, and fluoroquinolone was most prevalent (Fig. 2D). Indeed, the most abundant MDR ARGs in the infant gut were those that conferred resistance against both tetracycline and fluoroquinolone (Fig. 2D). We next investigated the abundance of MDR ARGs in different bacteria using MAGs (Fig. 2E). MDR ARGs were prevalent in infant gut bacteria, originating from 133 species and 8 phyla. Proteobacteria and Bacteroidetes had the highest abundance of MDR ARGs, with *E. coli* containing the most.

### Co-selection between ARGs and biocide or metal resistance genes is frequently detected in the infant gut, especially in *E. coli*

In environmental samples, it is well known that the persistence of ARGs is enhanced by co-localization with biocide resistance genes (BRGs) or metal resistance genes (MRGs). However, this phenomenon has yet to be fully characterized in humans, particularly in the early years of life. We therefore investigated in detail the co-localization between different types of resistance genes in the early gut, with respect to both co-resistance (ARGs and BRGs/MRGs with differing resistance profiles located in the same contig) and cross-resistance (a single gene conferring resistance to both antibiotics and biocides, ABRGs). Among all of the contigs that contained ARGs, 26.1% also carried BRGs that targeted a different mode of resistance (Fig. 3A). When we used MAGs to trace the bacterial origin of these contigs, we found that they originated from 5 phyla and 47 species; 92.2% of the traceable contigs carrying ARGs and BRGs with different resistance profiles came from Proteobacteria, of which 84.5% originated from *E. coli* (Fig. 3A). In the same way, we found that 15.5% of contigs with ARGs also carried MRGs (Fig. 3B), and 99.5% of the traceable co-localized contigs were of Proteobacterial origin, 91% from *E. coli* (Fig. 3B).

**Fig 3.**
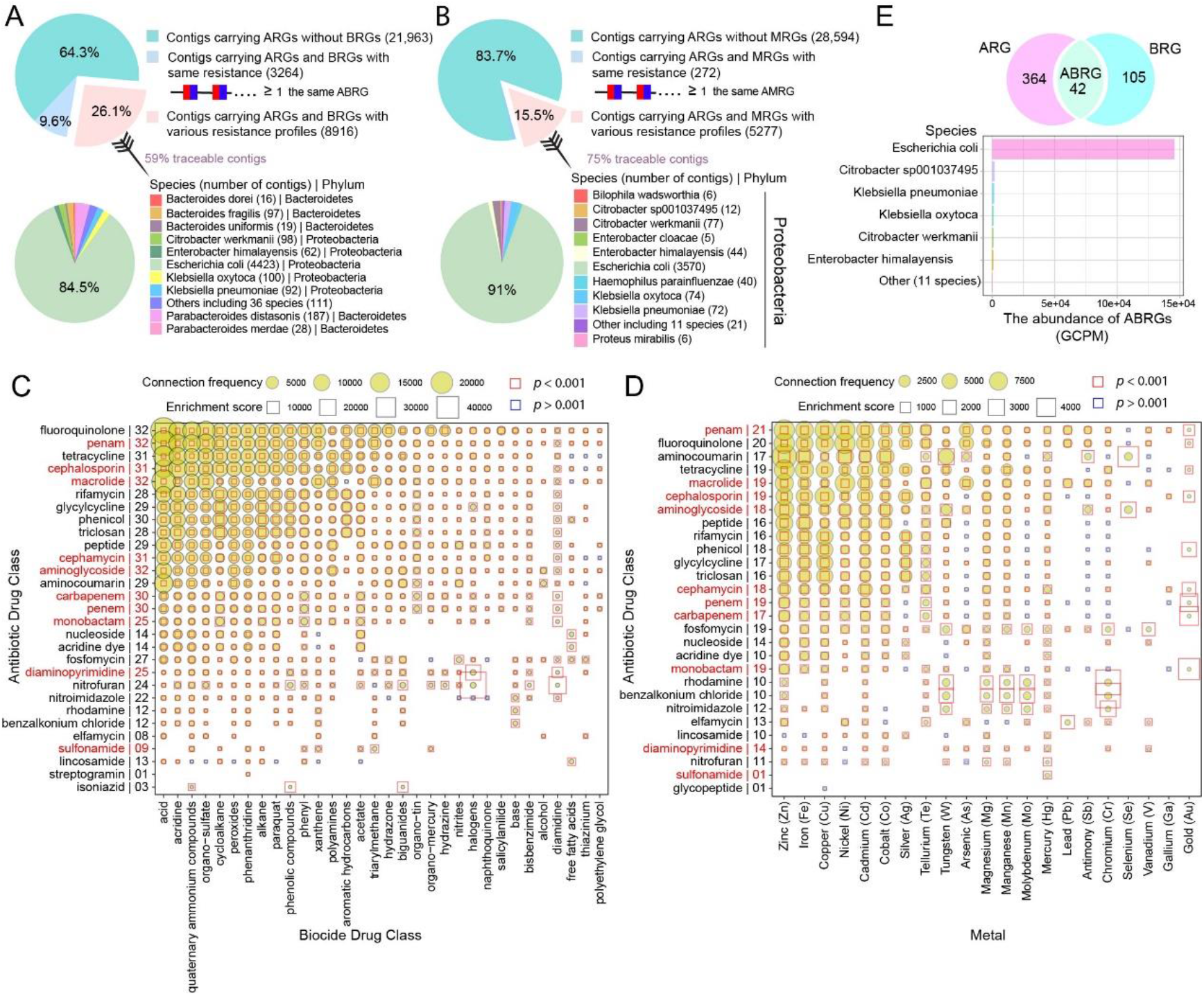
Co-localization between ARGs and BRGs or MRGs was frequently detected in the infant gut, especially in *E. coli*. **A)** Proportion of contigs carrying ARGs and BRGs, and the taxonomic origin (bacterial species and phylum) of contigs carrying ARGs and BRGs with different resistance profiles. **B)** Proportion of contigs carrying ARGs and MRGs, and taxonomic origin (bacterial species and phylum) of contigs carrying ARGs and MRGs with different resistance profiles. **C & D)** Co-localization bubble charts representing drug classes targeted by ARGs and BRGs with different resistance profiles (C) and between drug classes targeted by ARGs and metals targeted by MRGs (D). The frequency of connections in the contigs between ARGs and BRGs/MRGs with different targets, along with the enrichment scores, are shown in the figure. On the y-axis, the number to the right of the antibiotic drug class indicates the number of biocide drug classes (C) or metals (D) represented in the co-localization arrangements; the drug classes in red represent those used by members of this cohort. The size of a bubble indicates the frequency of connections in the contigs. A binomial test with FDR adjustment was used to test the statistical significance of enrichment patterns (p < 0.001 was set as significance cutoff; red square frame represents p < 0.001 and blue square frame represents p > 0.001). A significant p-value indicates that the occurrence of that specific pattern of gene co-localization would not be expected by chance. The size of the square frame indicates the magnitude of the enrichment score. **E)** Venn diagram showing the number of genes conferring antibiotic and/or biocide resistance and an abundance chart of ABRGs in different bacteria at the phylum and species level.

To better visualize gene co-resistance, we transformed the co-resistance network between ARGs and BRGs into a network representing 29 antibiotic drug classes and 32 biocide drug classes (Fig. 3C). The ARGs that most frequently co-occurred with BRGs were those that targeted fluoroquinolone, penam, tetracycline, cephalosporin, and macrolides. Moreover, ARGs associated with resistance to fluoroquinolone, penam, macrolides, and aminoglycoside were found to be co-localized with BRGs associated with all 32 biocides. The co-resistance network between ARGs and MRGs involved 28 antibiotic drug classes and 21 metals (Fig. 3D). The ARGs that most frequently co-occurred with MRGs were those conferring resistance to penam, fluoroquinolone, aminocoumarin, tetracycline, and macrolides; genes for penam resistance were found to be co-localized with resistance genes associated with all 21 metals. The co-localization enrichment scores were between 1.65 and 40,580 for combinations of ARGs and BRGs (mean (SD): 537(1814)) (Fig. 3C) and between 0.47 and 4,651 for ARGs and MRGs (mean (SD): 236(473)) (Fig. 3D). With the exception of ARGs related to lincosamide, most of the co-localization between ARGs and BRGs was significantly enriched compared to expected values (binomial test; adjusted *p* < 0.001, Fig. 3C), while 15% of the co-localization between drug resistance and metal resistance could have occurred by chance (binomial test; adjusted *p* >0.001, Fig. 3D). With respect to cross-resistance, approximately 10% of ARGs in the infant gut also conferred resistance to biocides, and these mainly originated from *E. coli* (Fig. 3E). The antibiotic and biocide resistance profiles that were most commonly implicated in cross-resistance were those related to fluoroquinolone and quaternary ammonium compounds (QACs), respectively (Fig. S1).

### Virulence genes are associated with an immature gut microbiome; co-localization is differentially abundant in different bacteria

Virulence factors can boost a bacterium’s ability to colonize and persist in the human gut by facilitating adhesion, nutrient competition, invasion, and immune defenses ^18^. Here, we detected a negative association between the abundance of virulence genes detected in the microbiome and gut microbial maturity at 1 year old (Fig. 4A). We assessed gut maturity using a ‘microbiota by age’ z-score (MAZ), with higher MAZ values representing higher maturity ^2^. Spearman correlation analysis revealed that a higher load of virulence genes was significantly correlated with lower MAZ scores (Fig. 4A, Spearman correlation, R = −0.27, *p* < 0.001), indicating that infants with a high load of virulence genes had more immature gut microbiomes.

**Fig 4.**
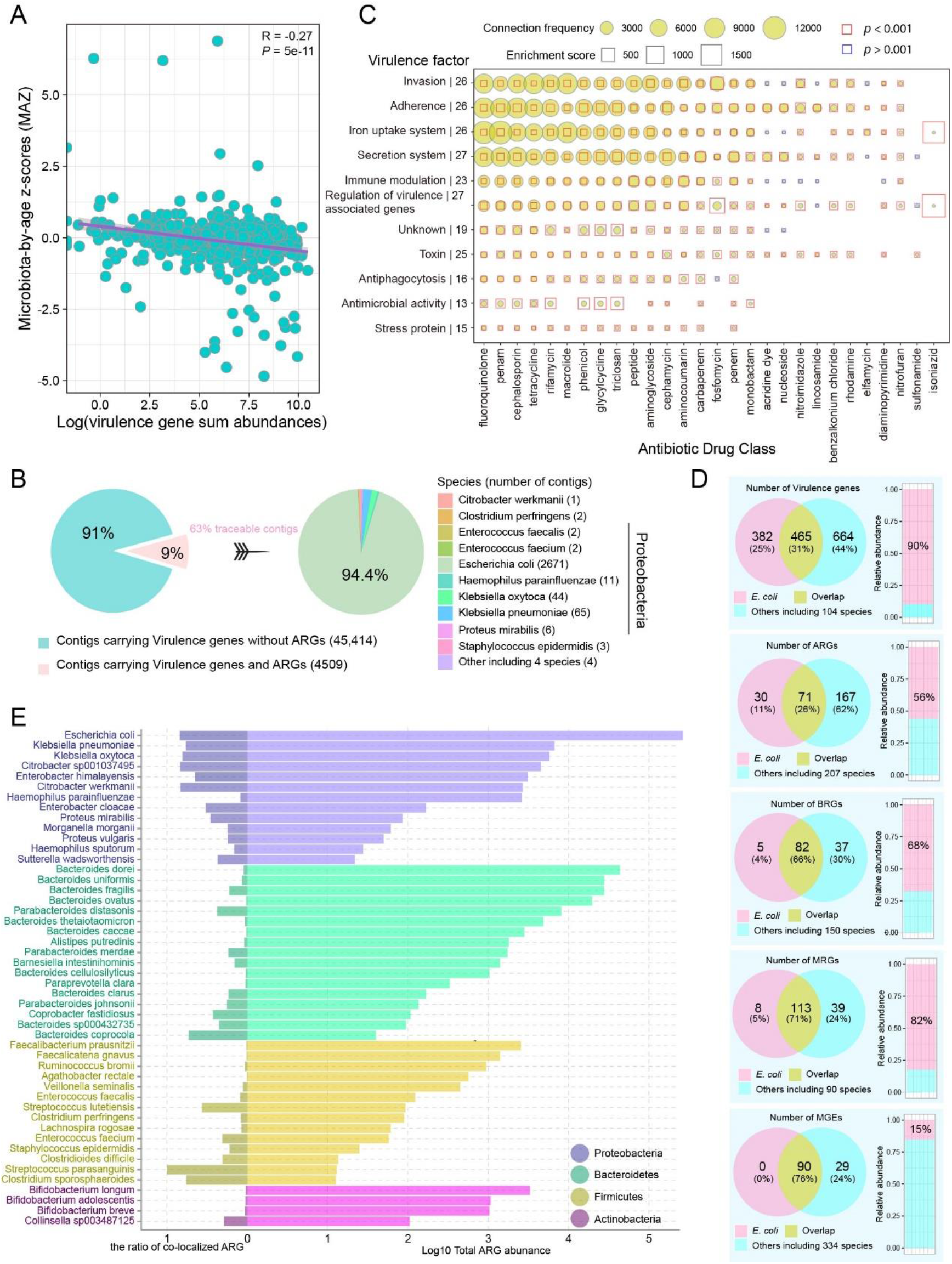
The association between virulence genes and gut microbial maturation, co-localization between ARGs and virulence genes, and differential patterns of co-localization among bacteria. **A)** The association between total abundance (log-transformed) of virulence genes and MAZ score. The confidence interval for the slope of the linear regression line is plotted as an illustration; inference was performed with the Spearman correlation coefficient R and the corresponding p-value (*p* < 0.05 as significance cutoff). **B)** Proportion of contigs carrying ARGs and virulence genes, and taxonomic origin (bacterial species and phylum) of contigs carrying both ARGs and virulence genes. **C)** Co-localization bubble chart depicting drug classes associated with ARGs and virulence factors encoded by virulence genes. The frequency of connections in the contigs between ARGs representing different drug classes and genes encoding different virulence factors are shown in the figure, along with enrichment scores. On the y-axis, the number to the right of each virulence factor indicates the number of antibiotic drug classes represented in the co-localization arrangements. The size of the bubble indicates the frequency of connections in the contigs. A binomial test with FDR adjustment was used to test the statistical significance of enrichment patterns (p < 0.001 was set as significance cutoff; red square frame represents p < 0.001 and blue square frame represents p > 0.001). A significant p-value indicates that the occurrence of that specific pattern of gene co-localization would not be expected by chance. The size of the square frame indicates the magnitude of the enrichment score. **D)** Venn diagram showing the number of virulence genes, ARGs, BRGs, MRGs, and MGEs in *E. coli* and other species and the relative abundance of these genes in *E. coli* compared to other species. **E)** The log-transformed total abundance of ARGs and the ratio of co-localized ARGs with BRGs in different bacterial species.

Co-localization between ARGs and virulence genes could enable the latter to gain a selective advantage from antibiotics and the former to gain enhanced persistence from increased colonization success. However, the prevalence of this phenomenon in the early gut is not clear. We therefore investigated the co-localization of ARGs and virulence genes in bacterial contigs in the infant gut; 9% of contigs with virulence genes also carried ARGs, with 94% of the traceable co-localized contigs originating from *E. coli* (Fig. 4B). The co-localization network of ARGs and virulence genes involved 28 drug classes and 11 virulence factors (Fig. 4C). Of particular interest were genes associated with virulence gene regulation and secretion systems, which were found to be co-localized with ARGs representing 27 different classes of antibiotics. Similarly, genes associated with invasion, adhesion, and iron uptake systems, also frequently appeared on the same contigs as ARGs. Enrichment scores for co-localization arrangements ranged from 0.98 to 1,943 (mean (SD): 97(177)) (Fig. 4C), and most of this enrichment was considered significant (binomial test; adjusted *p* < 0.001, Fig. 4C).

Overall, we found that co-localization between different types of resistance genes, or between resistance genes and those encoding virulence factors, was far more likely to occur in genomes in phylum Proteobacteria, especially in species in family *Enterobacteriaceae*, than in other bacteria. Proteobacteria, and *E. coli* in particular, appeared to serve as a reservoir of resistance genes and virulence genes in the infant gut (Fig. 4D, Fig. S2), which contributed to the bimodal distribution of these genes in this environment (Fig. S3). One potential explanation for a high frequency of co-localization could simply be high ARG content, i.e. species that contain more ARGs naturally host more co-localization. To test this hypothesis, we investigated patterns of co-localization between ARGs and BRGs. In the 50 bacterial species examined, we did not detect a significant positive correlation between the ratio of co-localized ARGs and total ARG abundance (Pearson correlation; R = −0.091, *p* = 0.53). This suggests that there is no uniform pattern of co-localization between ARGs and BRGs in all bacteria; instead, it appears that patterns of co-localization are specific to individual taxa. As shown in Fig. 4E, despite the large numbers of ARGs present in most species of Bacteroidetes, Firmicutes, and Actinobacteria, only a small number of these were found to be co-localized with BRGs. To further verify this disproportionate pattern of representation and exclude any confounding effect of differences in the lengths of assembled contigs in different phyla, we built a generalized linear model with a Poisson distribution to explore the association between the number of ARGs (the dependent variable), the presence of BRGs, and the log-transformed length of contigs in the four main phyla. Overall, Proteobacteria explained ≈29% of the variance in the co-localization of ARGs and BRGs, suggesting that this phylum was indeed the most important source for the co-localization of ARGs and BRGs.

### Antibiotics cause short-term abundance change in mobile ARGs; virulence genes exhibit higher potential for mobility than ARGs

Resistance and virulence genes can be widely transferred horizontally between bacteria via plasmids. We enumerated the distribution of different resistance genes on the plasmids (Fig. 5A). Approximately 1% to 9% of resistance genes and MGEs were located on plasmids (Fig. 5A). Many MGEs—such as transposons, integrons, and insertion sequences—can transfer genetic elements between intracellular chromosomes and plasmids, or between different plasmids ^19^, and often appear next to the transferred genes. We thus explored mobile co-localization phenomena on plasmids involving the following elements (Fig. 5B): 1) ARGs, BRGs, and MGEs; 2) ARGs and MGEs (referred to as mobile ARGs); and 3) virulence genes and MGEs (referred to as mobile virulence genes).

**Fig 5.**
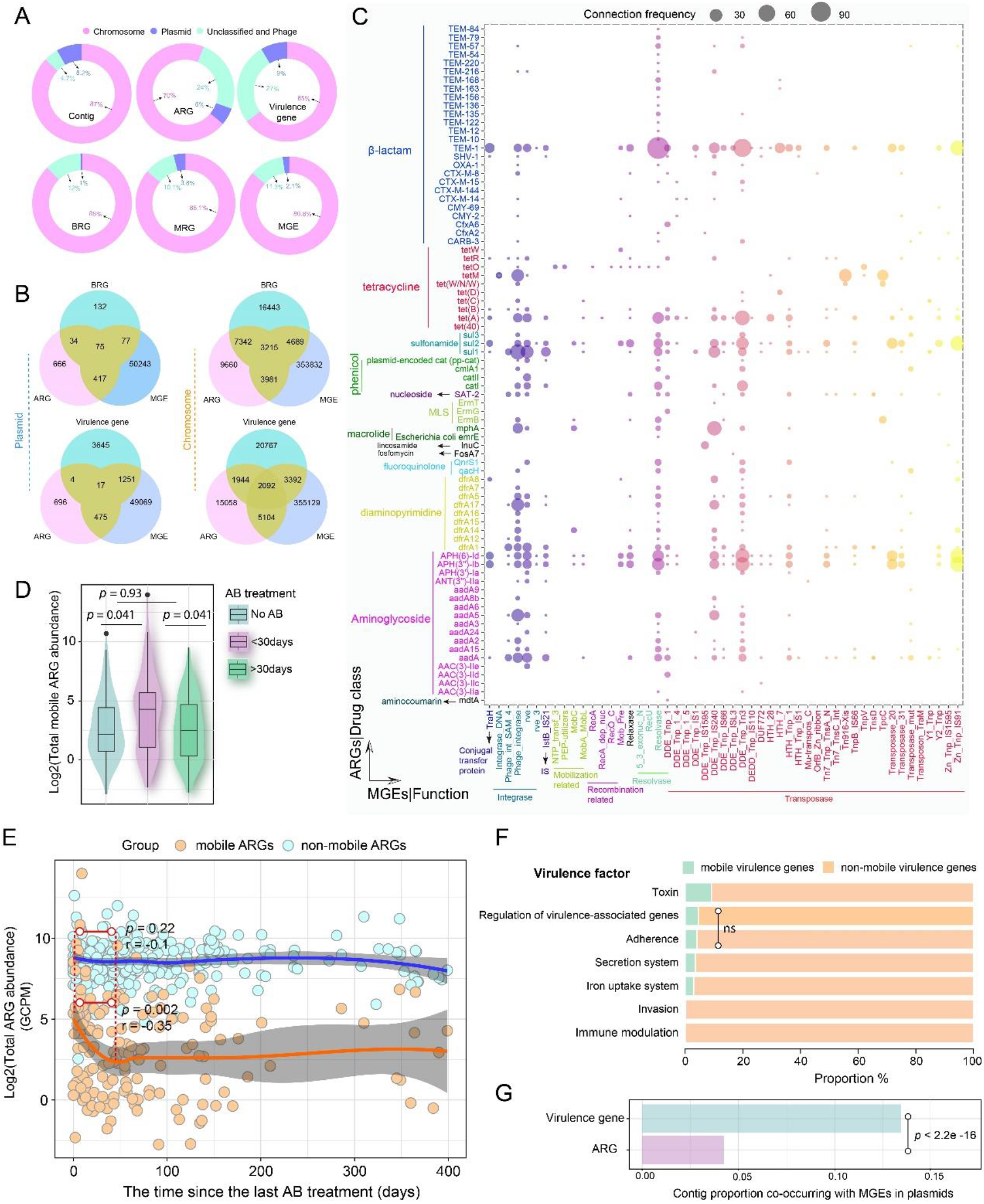
Co-localization of ARGs, BRGs/virulence genes, and MGEs on plasmids in the infant gut. **A)** The respective abundance (percentage) of contigs, resistance genes, virulence genes, and mobile genetic elements in chromosomes and plasmids. **B)** Venn diagram depicting the number of contigs with resistance genes and MGEs in chromosomes and plasmids. **C)** Co-localization bubble chart representing ARGs and MGEs in plasmid contigs. The size of the bubble is proportional to the frequency of connections in the contigs. **D)** Log-transformed total abundance of mobile ARGs in the gut of infants who had taken antibiotics < 1 month before sampling, > 1 month before sampling, or not at all. FDR-adjusted *p* values listed in the figure were calculated with pairwise Wilcoxon rank-sum tests. **E)**The effects of antibiotic treatment on log-transformed total abundance of mobile ARGs and non-mobile ARGs with time. Confidence interval for the slope of the linear regression line is shown. The significance level and Pearson correlation coefficient R are shown in the figure; dotted lines denote the window in which the Pearson correlation was calculated (from 0 to 45 days). AB represents antibiotics. **F)** The relative proportions of mobile and non-mobile virulence genes related to seven virulence factors. Pairwise Fisher’s exact test was carried out for the comparison of the relative proportions of mobile and non-mobile genes within each of the seven groups. Multiple comparisons were adjusted for FDR. Except for the pairwise comparison of adherence- and regulation-associated genes (*p* = 0.06, indicated by ns), there were significant differences between all paired comparisons (*p* < 0.001). **G)** The proportion of all contigs with ARGs/virulence genes and MGEs that were found in plasmids. *P*-value from Fisher’s exact test is shown in the figure.

Within the infant gut, class 1 integrons were the most common co-localization loci residing on plasmids. We detected 36 representative plasmid contigs carrying ARGs, BRGs, and MGEs. Of these, 22 contained a complete class 1 integron (Fig. S4), featuring a 3’-conserved segment containing a sulphonamide resistance gene (*sul*) and a QAC resistance gene (*qac*), a 5’-conserved segment carrying an integrase, and a gene cassette. In total we found that 17 ARGs were integrated into the cassette (Fig. S4), including six aminoglycoside resistance genes (*aadA*/*2*/*3*/*5*/*6*/*8b*,) seven diaminopyrimidine resistance genes (*dfrA1*/*5*/*7*/*12*/*15*/*16*/*17*), two sulfadiazine resistance genes (*sul1*/*3*), one beta-lactamase resistance gene (*CARB*-*3*), and one phenicol resistance gene (*cmlA1*).

We identified 80 mobile ARGs that co-localized with 51 MGEs on plasmids (Fig. 5C). Of the mobile ARGs, 26 encoded resistance to β-lactams, 17 to aminoglycoside, and 10 to tetracycline. Among the different MGEs, transposases and integrases were most commonly found beside ARGs on plasmids. When we examined the influence of antibiotic use on the abundance of mobile ARGs in the infant gut, we found a transient effect: the total abundance of mobile ARGs in the gut was lower in infants who had received antibiotic treatment within 30 days prior to sampling than in those who had received antibiotic treatment more than 30 days before sampling or not at all. Instead, no significant differences were found between the latter two groups (Fig. 5D). Furthermore, in infants who received antibiotics, the total abundance of mobile ARGs in the gut peaked on the first day after treatment (Fig. 5E), significantly declined over the next ca. 45 days (Pearson correlation, R = −0.35, *p* = 0.002), and finally reached a stable plateau close to the levels found in the guts of infants who had not received any antibiotic treatment. In contrast, putatively non-mobile ARGs, i.e. those not on the same contigs as MGEs, did not exhibit any change in abundance in the 45 days after antibiotic usage (R = −0.1, *p* = 0.22). These results indicate that antibiotic treatment exerted an influence mainly on mobile ARGs.

Although HGT is thought to be critical for the transmission of virulence genes in the early gut microbiota, it has been poorly characterized in the infant gut. Among the 2128 total virulence genes in our sample, we detected 321 mobile virulence genes; the majority of these were associated with adhesion, regulation, secretion systems, toxins, and iron uptake systems (Fig. 5F and Table S1). Genes associated with toxins showed the strongest mobility, followed by those related to regulation, adherence, secretion, iron uptake, invasion, and immune modulation. We detected significant differences in the proportions of mobile genes among virulence categories (pairwise Fisher’s test with FDR adjustment, *p* < 0.001 for each comparison, Fig. 5F), with the exception of genes related to adhesion and regulation (*p* = 0.06, Fig. 5F). Among the contigs that contained mobile virulence genes (Fig. 5B), 5.3% carried over 10 virulence genes and could thus represent pathogenicity islands. In particular, when we compared contigs carrying mobile virulence genes with those carrying mobile ARGs, we observed that a significantly higher proportion of the former originated from plasmids (13.4% of contigs with mobile virulence genes; N = 1,268) than of the latter (4.2% of contigs with mobile ARGs; N = 492) (Fisher’s exact test, *p* < 0.001, odds ratio = 3.2 (95% CI: 2.84–3.53)) (Fig. 5G). To exclude the confounding influence of assembled contig lengths, we built a logistic regression with a binomial distribution to explore the association between the presence of virulence genes, ARGs, and MGEs in contigs (the dependent variable), the presence of these contigs in plasmids and the log-transformed length of these contigs. Even after adjusting for contig length, we found that significantly more virulence genes were present in plasmids (odds ratio= 1.88 (95% CI: 1.74–2.04), *p* < 0.001).

### The abundance of co-localized ARGs associates with infant gut microbial maturity

High ARG abundance is linked to low gut microbial maturity in infants ^6^, and the co-localization of ARGs may promote the persistence of this state in the infant gut. To shed light on this phenomenon, we explored the relationship between the abundance of co-localized ARGs and gut microbial maturity at one year of age, as quantified by a ‘microbiota-by-age’ z-score (MAZ) calculated with a previously developed model ^2^. Linear regression analysis revealed that a higher abundance of co-localized ARGs was significantly correlated with lower MAZ scores (Fig. 6A), i.e. immaturity. Maturation of the gut microbiome is a complex process and represents shifts in the composition of bacterial communities with age ^2^. However, given the vital role played by *E. coli* in providing co-localized ARGs to the microbial community, it is possible that the relationship between co-localized ARG abundance and gut maturity may depend solely on the abundance of *E. coli*. To investigate this hypothesis, we first fit a linear regression model with the log-transformed relative abundance of *E. coli* as the explanatory variable and MAZ score as the dependent variable, and found a significant relationship between the two (Fig. 6B, estimate −0.16 SD per log10 fold change, 95% CI [−0.23, −0.08], *p* < 0.001). This suggested that the high abundance of *E. coli* in the gut of one-year-old infants was associated with low gut microbial maturity. Using 16S sequencing data from different time points, we found that the mean abundance of *E. coli* peaked at birth and then gradually decreased, reaching relative equilibrium after 3–4 years of age (Fig. 6C). When we added the abundance of co-localized ARGs and BRGs to the model (Fig. 6E), we found that this was also associated with the MAZ score (*p =* 0.02) and, in fact, altered the relationship between *E. coli* and MAZ (*p* = 0.21). Likewise, when we added the abundance of co-localized ARGs and MRGs or virulence genes to the model, there was also a significant effect on MAZ (*p* = 0.0005, 0.0002) that changed the relationship between *E. coli* and MAZ (*p* = 0.19, 0.42) (Fig. 6E). Thus, after accounting for the abundance of co-localization in these cases, the abundance of *E. coli* was no longer a significant factor in the models. This suggests that most of the association between *E. coli* and gut maturity which was captured by its ARG co-localized gene abundance is not a sole variable in terms of their association with maturity.

**Fig 6.**
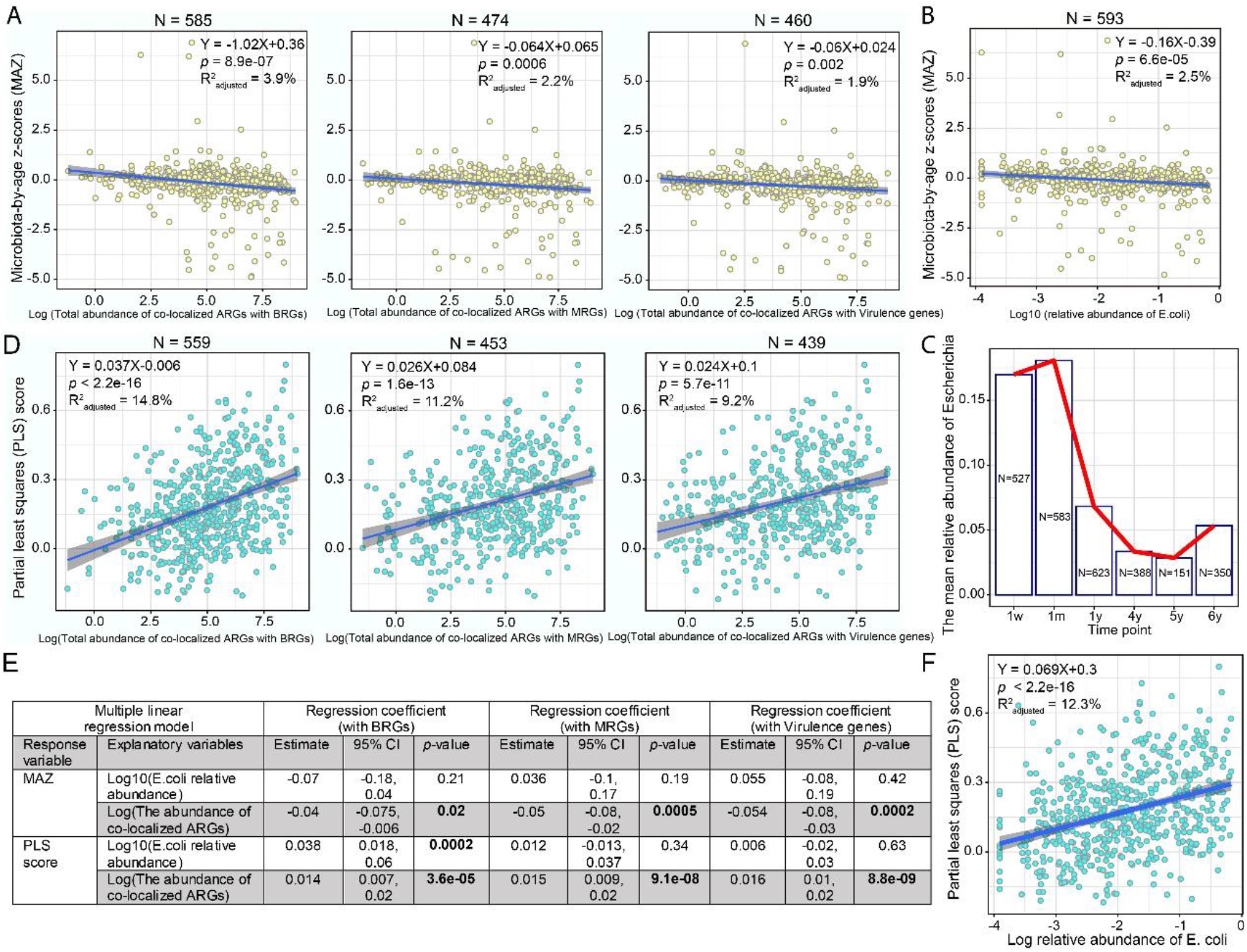
The association between the abundance of co-localized ARGs and infant gut microbial maturity/asthma-associated microbiome. The association between log-transformed total abundance of co-localized ARGs **(A, D)** or *E. coli* relative abundance **(B, F)** and MAZ/PLS score. Confidence interval for the slope of the linear regression line is shown. The formula of the regression line, *p*-value for slope, and adjusted R^2^ value are listed in the figures. The number of samples used in the linear regression analysis is denoted by N. **C)** The mean relative abundance of *E. coli* in the gut of infants at one week, one month, one year, four years, five years, and six years of age. **E)** Multiple regression analysis evaluating whether *E. coli* is the sole determinant of the relationship between co-localized ARGs and infant gut microbial maturity/asthma-associated bacterial community.

In previous work, we constructed a cross-validated sparse partial least squares (PLS) model to investigate the association between gut bacterial composition at one year of age and asthma at five years of age ^2^. This model was based on bacterial abundance from 16S sequencing data and thoroughly validated asthma phenotypes in the COPSAC2010 birth cohort. Higher PLS scores for gut bacteria at age one predicted a higher risk of developing asthma at age five. Here, we observed a positive correlation between the abundance of co-localized ARGs and asthma-associated bacterial PLS scores (Fig. 6D), implying a degree of similarity between a gut microbial community that is rich in co-localized ARGs and one that is associated with asthma. Considering the dominant role of *E. coli* in ARG co-localization, we next verified whether this correlation was only related to *E. coli* abundance. In a univariate linear model, *E. coli* abundance was positively correlated with PLS score (Fig. 6F). When the model was modified to include the abundance of ARGs co-localized with BRGs, MRGs, or virulence genes (Fig. 6E), these three factors also correlated with PLS score (*p* < 0.0001). However, the latter two altered the association between *E. coli* and PLS so that it was no longer significant (*p* = 0.34, 0.63). These results suggest that the abundance of ARG co-localization and *E. coli* shared a large amount of information on the asthma-associated microbial community. It appears that a gut microbial community that features an abundance of co-localized ARGs is somehow associated with an increased risk of later asthma, but the exact mechanism remains unclear.

## Discussion

In this study, we comprehensively analyzed genetic co-localization involving ARGs and the association of this phenomenon with environmental factors and gut microbial maturation in a cohort of 662 Danish children. Collectively, the most common forms of co- and cross-resistance in the infant gut microbiome were associated with the broad-spectrum antibiotics fluoroquinolone and tetracycline, and derived largely from *E. coli*. Multidrug resistance against fluoroquinolones and tetracycline has also been reported to be prevalent in ESBL producers (e.g., *E. coli*) isolated from multiple ecological settings, including water bodies and the feces of animals and healthy humans ^20–22^. This pattern is consistent with the extensive use of these antibiotics in animal husbandry and medicine in recent decades, and also involves the HGT of resistance genes on MGEs ^23–28^. In addition, with the extensive use of other antimicrobial agents, co-selection of resistance to antibiotics, biocides, and/or metals has been commonly detected in bacteria from the environment, animal farming, and the human microbiome ^28–34^. Similarly, we also observed ARGs and BRGs or MRGs co-occurring on the same contigs in the infant gut at frequencies higher than those expected by chance, suggesting an enrichment effect. However, the extent to which antimicrobial agents and MGEs drive this enrichment effect, i.e. the mechanism of co-selection and the underlying drivers, is not entirely clear ^35^. Multi-resistant bacteria such as *E. coli* are some of the first colonizers in the gut at the beginning of life ^6,36^, which means that co-selection between different types of antimicrobial resistance has become a critical One Health issue ^37^.

Co- and cross-resistance to antibiotics and biocides, along with multiple-antibiotic resistance, were primarily found in Proteobacteria, especially in the clinically relevant bacterial family *Enterobacteriaceae*. Members of this family, especially *E. coli* and *Klebsiella pneumoniae*, are among the most important causes of serious nosocomial and community-acquired bacterial infections in humans, and frequently exhibit drug resistance ^38^. The abundant multidrug pumps in *Enterobacteriaceae*^39–41^ can actively expel multiple antibiotics and biocides from the cell ^42^ and are therefore the main contributor to cross-resistance between different drugs. The high prevalence of co-localization in family *Enterobacteriaceae* was also observed in previous research ^34,43^.

The presence of virulence genes together with ARGs may result in the emergence of novel “superbugs”^44^—pathogenic bacteria with multi-resistant phenotypes. A recent large genomics investigation confirmed abundant co-localization between virulence genes and ARGs in human pathogenic bacteria, and this co-occurrence or correlation was observed across various ecological settings ^45–48^. Similarly, our study identified abundant co-localization in opportunistic pathogens such as *E. coli*, *H. parainfluenzae*, *Klebsiella spp*., *Enterococcus spp*., and *Citrobacter spp*. in infants. Through HGT of plasmids, multiple co-localized resistance genes can easily spread among bacteria. In the plasmids examined in this study, class 1 integrons were identified as the predominant genetic loci for the transfer of ARGs with BRGs and/or MRGs in the infant gut. This is consistent with previous work reporting that most known ARG cassettes are located in class 1 integrons ^49^, which have been identified in more than 70 clinical bacterial species and are affected by human activities ^50^. Over the last few decades, QACs have been widely used as cationic surfactants in household products, and this has been accompanied by a concomitant increase in QAC resistance genes in bacteria, thereby increasing the risk of co-selection of antibiotic-resistant bacteria ^51–53^. Accordingly, our study reveals that co-selection of QAC resistance genes and ARGs has occurred in healthy infants.

In addition, we explored mobile ARGs and virulence genes in the infant gut. Our results demonstrate that antibiotic treatment caused a short-term change in the total abundance of mobile ARGs in the infant gut, suggesting that antibiotic use facilitated the acute spread of resistance among bacteria, with MGEs as vectors. A similar association between oral antibiotic use and increased intestinal MGE abundance was reported in a study on the gut microbiome of fish ^54^. Here, we found a significantly higher proportion of mobile virulence genes than mobile antibiotic resistance genes. The transfer of virulence genes is one of the main ways in which bacterial pathogens acquire virulence in the course of evolution ^17^, and, as reported here, a variety of virulence genes, including adherence factors, secretion systems, and toxins, have been detected in plasmids ^55–57^. HGT events involving virulence genes pose a particular threat to health through, for example, the emergence of novel pathogens ^58^ and the formation of defense islands or pathogenicity islands over time ^59^.

The abundance of co-localized ARGs significantly influenced the maturation of the infant gut microbiome, with *E. coli* playing a vital role. As one of the earliest colonizers, the abundant resistance and virulence genes in *E. coli* (Fig. S2) can grant it a substantial selective advantage. However, the prolonged persistence of *E. coli* may disrupt the subsequent colonization of beneficial commensal bacteria, thereby leading to a delay in the maturation of the infant gut microbiome. Early intervention is therefore essential. However, due to the high genetic variability of *E. coli*^60^, its persistence and harmfulness to the host may vary considerably among strains ^61,62^. Future research focused on differentiating the effects of different *E. coli* strains on gut microbial maturation would be useful in identifying the strains for which intervention is most critical.

In conclusion, we found that the phenomenon of co-localization between ARGs and other resistance and virulence genes was prevalent in the gut at the beginning of life. This is a critical One Health issue that emphasizes the need to apply caution in the use of antimicrobial agents in clinical practice, animal husbandry, and daily life. The evidence presented here not only describes ARG colonization early in life, but also suggests the means by which resistance is maintained in the gut, potentially leading to long-term health threats such as delayed gut maturation in infants and its associated downstream consequences. Our study provides a thorough characterization of the genetic basis for drug resistance in infants and forms the basis for further investigation into the clinical use of pediatric antibiotics.

## Methods

### Study population and sample collection

The study subjects were participants in the population-based COPSAC2010 mother-child cohort consisting of 700 mother-child pairs ^63,64^. The fecal samples investigated in this study were gathered from infants at 1 year of age, either at the COPSAC research unit or by the parents at home, following instructions. Each collected fecal sample was blended with 1 ml of 10% glycerol and stored at −80°C for storage before DNA extraction.

### Ethics

The study was approved by the National Committee on Health Research Ethics (H-B-2008-093) and the Danish Data Protection Agency (2015-41-3696), and was supervised according to the criteria set by the Declaration of Helsinki. For each child, both parents provided their oral and written informed consent before enrollment.

### Metagenomics sequencing and data processing for fecal samples

Bacterial DNA from 663 fecal samples was extracted using the PowerMag®Soil DNA Isolation Kit optimized for the epMotion robotic platform model according to extraction instructions. The Kapa Hyper Prep kit (for Illumina) was used for sequencing library preparation. Fecal DNA samples were sequenced using the Illumina NovaSeq apparatus by Admera Health (USA). Out of the 663 samples, one sample failed to produce a library. To avoid a batch effect, the 663 samples were sequenced in the same batch. GNU Parallel v20180722^65^ was used for parallelized preprocessing during bioinformatics analysis. The adapter sequences were trimmed by BBDuk (BBTools v38.19) using the default setting except for the following parameters: “ktrim=r k=23 mink= 11 hdist=1 hdist2=0 ptpe tbo”. Reads shorter than 50 bases and low-quality sequences were removed by Sickle v1.33^66^. Human genome contaminants were filtered out using BBMap (BBTools v38.19) with the default setting. Short reads were assembled into contigs individually using SPAdes v3.12.0 under the default settings ^67^. Metagenomic sequencing coverage was analyzed using Nonpareil v3.30 with kmer mode ^68^. Taxonomic classification of microbial communities was inferred with humann2 v0.11.2^69^ and MetaPhlAn v2.7.5^70^. The binning of metagenomics contigs into metagenomically assembled genomes (MAGs) individually was performed by three binners in the metaWRAP pipeline (v1.2.2)^71^: MetaBAT2 v2.12.1^72^, MaxBin2 v2.2.6^73^, and Concoct v1.0.0^74^. The quality assessment of MAGs was carried out using CheckM v1.0.12^75^, and only MAGs with at least 90% integrity and no more than 5% contamination were retained. One sample did not generate any MAGs. In total, 452 dereplicated MAGs were generated for the 661 samples. GTDB-Tk toolkit (v1.7.0 and GTDB-Tk reference data r202) was used to infer the bacterial taxonomic assignments of MAGs ^76^. Open reading frames (ORFs) in contigs were identified with Prodigal v2.6.3 in META mode ^77^.

### Identification of resistance genes, MGEs, virulence genes, and integrons

Resistance Gene Identifier ^78^ was used to search for ARGs within the predicted ORFs by aligning the amino acid sequences of ORFs to the Comprehensive Antibiotic Resistance Database (CARD v3.0.7). The significance cut-offs “Perfect” (100% identity and 100% reference sequence coverage) and “Strict” (a match higher than the bitscore of the curated BLASTP bitscore cutoff) were used as the thresholds for filtering ARGs. Diamond blastp was used to search for BRGs and MRGs in the predicted ORFs by aligning the amino acid sequences of ORFs to sequences in a database of antibacterial biocide and metal resistance genes (BacMet v2.0)^79^ using the more sensitive mode and k1 option ^80^. Thresholds of 90% identity and 80% query coverage were used to predict BRGs and MRGs. MGE homologs were characterized using the PFAM ^81^ and TnpPred ^82^ databases through HMMSEARCH (v3.1b2) ^29,83^, with “-cut_ga” as the threshold. If ORFs had multiple hits, we only kept the one with the lowest E-value. Diamond blastp search was also performed to predict the families of bacterial virulence factors from the amino acid sequences of the predicted ORF using the VFDB database (updated version on Sep 10, 2021)^84^ with the more sensitive mode and k1 option ^80^. Thresholds of 90% identity and 80% query coverage were used to predict virulence genes. The profiles of resistance genes and virulence genes are shown in Fig. S5. *AttC* recombination sites, promoters, and *attI* sites for the integrons were identified by IntegronFinder with the default setting ^85^.

### Calculation of gene abundance

The alignment of clean reads against the ORFs was performed by Bowtie2 aligner ^86^. Samtools idxstats ^87^ was used to calculate the number of mapped reads in the bam file. We used values of gene coverage per million (GCPM) for ORFs to normalize the length of ORFs and sequencing depth ^29^. Because the sum of GCPM values of all the ORFs in each sample is the same, the abundance of ORFs is thus comparable between samples. The formula to calculate GCPM is 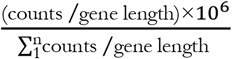, where counts represents the number of mapped reads, gene length represents the ORF length, and n represents the total number of predicted ORFs in each sample.

### Identification of bacterial origin of genes

The bacterial species from which gene-containing chromosomal contigs originated were traced from the taxonomic classification of MAGs. In this way, we were able to classify genes to their bacterial species of origin.

### Co-localization analysis

Co-localization represents the physical relationship between genes on the same assembly contig. We selected representative plasmid contigs to demonstrate class 1 integrons; first, we deduplicated co-localization contigs based on gene combinations, then removed the co-localization contigs whose gene combinations were included in other co-localization contigs and the representation contigs were finally obtained. Gene arrangements in the representative contigs were visualized using gggenes (https://wilkox.org/gggenes/). In this study, we refer to co-localization associated with MGEs occurring on plasmids as mobile co-localization. The three types of mobile co-localization phenomena investigated here are 1) ARGs, BRGs, and MGEs; 2) ARGs and MGEs; and 3) virulence genes and MGEs. The ARGs and virulence genes in categories 2 and 3 are also referred to as mobile ARGs and mobile virulence genes.

### Source identification of co-localization contigs

We used PPR-Meta v1.1^88^ to classify metagenomics sequences as coming from chromosomes, plasmids, or phages. As the software suggested, a probability score of 0.7 (between 0 and 1) was used as the threshold. The bacterial species of origin of chromosomal contigs was traced from the taxonomic classification of MAGs.

### Enrichment score

We assessed patterns of co-localization using an enrichment score, which we defined as the fold difference between the actual and the expected numbers of co-localization contigs. This was calculated as 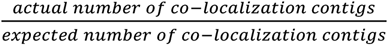. A score higher than 1 served as evidence of enrichment in the gut. A binomial test with FDR adjustment ^89^ (R function “binom.test”) was used to test whether the actual number of co-localization contigs was significantly higher than the expected number of co-localization contigs (*p* < 0.001 as significance cutoff), i.e. whether an enrichment score was statistically significant and a given pattern of gene co-localization occur by chance. The x, n, and p in the R function “binom.test” represent the actual number of co-localization contigs carrying resistance genes for drugs A and B, the total number of contigs, and the expected probability of contigs carrying resistance genes for drugs A and B, respectively. The expected probability p calculated as 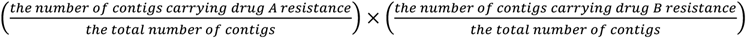.

### Statistical Analysis

The statistical software “R” was utilized for data organization and statistical analyses ^90^.

#### Gene profile clustering analysis

The ‘partitioning around medoids’ (PAM) clustering approach ^91^ (R-package “cluster”^92^) was used to cluster BRGs/MRGs/virulence genes. The Euclidean distance between clusters was used to estimate the average silhouette width (higher average silhouette width indicates better clustering). The number of gene clusters with the highest average silhouette width was the optimal cluster number (R-package “fpc”^93^).

#### Differential abundance analysis

Differences in gene abundance between two groups were compared by pairwise Wilcoxon test. All tests were adjusted for multiple comparisons using the FDR algorithm ^94^.

#### Modeling analysis of co-localization between ARGs and BRGs among different bacteria

To check for differential patterns of co-localization between ARGs and BRGs in different bacteria and adjust for the different lengths of co-localized contigs, we built a generalized linear model with a Poisson distribution using R function “glm”, and used a Poisson regression to evaluate the association between the numbers of co-localized ARGs (dependent variable), the presence of BRGs, and the log-transformed length of co-localized contigs in the four main bacterial phyla.

#### Modeling analysis of mobile ARGs and virulence genes in plasmid contigs

We built a logistic regression with a binomial distribution using R function “glm” and used a binomial regression to explore the association between the presence of mobile virulence genes or ARGs (i.e. virulence genes or ARGs with MGEs) in contigs (dependent variable), the presence of these contigs in plasmids, and the log-transformed length of these contigs.

#### Correlations between gut microbial maturity/later asthma risk and the abundance of co-localized ARGs

In previous work, we demonstrated how to calculate a microbiota-by-age z-score (MAZ) and sparse partial least squares (PLS) score for evaluating gut microbiome maturity and asthma risk at age 5, respectively ^2^. Here, we created a linear model that evaluated the relationship between PLS score/MAZ and the abundance of *E. coli* and co-localized ARGs.

#### Bacterial ranking by random forest analysis

The importance of individual bacterial species in shaping the gene clusters was evaluated by Random Forest analysis ^95^ (R-package “randomForest”^96^). In this model, the number of variables per split and the number of trees were set to 50 and 500, respectively, resulting in a steady classifier with a low error rate (≈7%). The importance of a given variable was evaluated by the mean decrease in accuracy (a higher value means a more important variable).

### Sequencing of 16S rRNA gene amplicons from fecal samples at different time points

This work is described in more detail in a previous paper ^6^. Briefly, a total of 2622 longitudinal stool samples were collected at the ages of one week, one month, one year, four years, and five years old (N = 527, 583, 623, 388, 151 and 350, respectively). The V4 region of the 16S rRNA gene was sequenced for all samples. The DADA2 QIIME 2 platform was used to characterize ASVs and perform taxonomic assignments across samples ^97,98^.

## Data accessibility

All sequencing data are available in the Sequence Read Archive (SRA) under the accession number PRJNA715601. According to Danish and European law, data that involve the personal privacy of project participants cannot be publicly available without a cooperation agreement and a data transfer agreement.

## Code availability

The R code for data analysis is available upon request from the authors.

## Acknowledgments and funding

We are grateful to the children and families who participated in the COPSAC2010 cohort study for their dedication and support. Furthermore, we acknowledge and appreciate the unique efforts of all members of the COPSAC research team. COPSAC is supported by a variety of private and public research funds, all of which can be found at www.copsac.com.

